# Local two-photon excitation reveals the mechanism of mitochondrial Ca^2+^ release

**DOI:** 10.1101/2023.08.31.555665

**Authors:** Bingyi Li, Xiaoying Tian, Shaoyang Wang, Yujie Zhu, Hao He

## Abstract

Mitochondrial Ca^2+^ (mitoCa^2+^) simultaneously implicates respiration, mitochondrial physiology, and cell signaling, which prevents the disentanglement of mitoCa^2+^ from those complex processes. Although mitochondria have long been recognized as temporary Ca^2+^ buffer, how mitoCa^2+^ is regulated and released remains unclear. In this study, we report a specific photochemical excitation to flavoproteins in single-mitochondrion tubulars by a tightly-focused femtosecond laser that triggers local mitoCa^2+^ transients, without any extra-mitochondrial Ca^2+^ involved. The two-photon excitation to complex I and II accelerates the entire electron transport chain (ETC) and promotes ATP synthesis. Free mitoCa^2+^ is thus released from the Ca^2+^-phosphate ion (Pi) complexes at complex V in mitochondrial matrix during ATP synthesis to form mitoCa^2+^ transients there. The abnormal mitoCa^2+^ signaling by knockdown of ATP synthase subunit affects cell proliferation, apoptosis, and mitophagy. Our results reveal mitoCa^2+^ is released and regulated by ETC and ATP synthesis rather than the reverse.

Mitochondria are multi-functional units to simultaneously produce energy and coordinate molecular signaling for cells. The key process of metabolism, tricarboxylic acid (TCA) cycle, is continuously ongoing in mitochondrial matrix and generates reducing equivalents for subsequent electron transfer (Martinez-Reyes and Chandel, 2020; Martinez-Reyes et al., 2016). The respiration is accomplished by electron transport chain (ETC) in the inner mitochondrial membrane (IMM) (Marreiros et al., 2016). Mitochondria regulate a series of signaling cascades to mediate apoptosis, autophagy, and cell senescence (Galluzzi et al., 2014; Rasola and Bernardi, 2011; Ziegler et al., 2021). Biosynthesis of some amino acids and nucleotides also takes place in mitochondria (Ahn and Metallo, 2015; Li and Hoppe, 2023). Those processes are involved with each other through complex crosstalk and feedbacks.

Mitochondria have long been recognized as Ca^2+^ buffer to temporarily deposit abnormal cytosolic Ca^2+^ for cellular Ca^2+^ homeostasis (Garbincius and Elrod, 2022; Kirichok et al., 2004; Lambert et al., 2019). However, the Ca^2+^ entry into mitochondria definitely influence those mitochondrial processes and functions (Garbincius and Elrod, 2022). The acute and direct consequences of Ca^2+^ entry into mitochondria are found as depolarization of mitochondrial membrane potential (MMP). If the cellular Ca^2+^ level is too high, mitochondria may become damaged and dysfunctional. Mitophagy/autophagy and even apoptosis are initiated (Galluzzi et al., 2014; Lou et al., 2020; Rasola and Bernardi, 2011). Moreover, Ca^2+^ in mitochondrial matrix (mitoCa^2+^) takes an essential role in quite a few physiological processes including mitochondrial fission and fusion, cell development, and proliferation (Doonan et al., 2014; Singh and Mabalirajan, 2021; Steffen and Koehler, 2018). The respiration is also believed to be related to mitoCa^2+^ (Glancy and Balaban, 2012; Wescott et al., 2019). Technically, perturbation to mitoCa^2+^ affects most mitochondrial processes and physiology, which makes it quite difficult to interrogate how Ca^2+^ is regulated and released in mitochondria. So far, the regulation of mitoCa^2+^ remains in mist.

In this study, we report a single-mitochondrion photochemical process by tightly-focused femtosecond laser that specifically excites complex I and II by two-photon excitation and accelerate ETC. Free mitoCa^2+^ is released from Ca^2+^-phosphate ion (Pi) complexes by ATP synthesis at complex V in mitochondria. These results clarify the mechanism of mitoCa^2+^ regulation and provide further insights in the relationship between mitoCa^2+^ and respiration.

## Results

### 1. ETC acceleration triggers local mitoCa^2+^

We at first reported a two-photon photochemical excitation to flavoproteins that further triggers local mitoCa^2+^ transients at single-mitochondrion level. This process could be accomplished on any two-photon or confocal (coupled with a femtosecond laser) microscope systems. With mitoCa^2+^ fluorescently indicated, an immediate and dramatic local mitoCa^2+^ rise in a single mitochondrial tubular structure could be activated if it was transiently excited by a tightly-focused femtosecond laser at 700 nm for 0.1 s at 5 mW (Fig. 1 A). In this scheme, the laser was focused to a submicron-level diffraction-limit spot (∼ 0.6 μm in diameter, focused by a ×60 objective, N.A. = 1.2), which ensured precisely locating the photoexcitation target to a single tubular of mitochondrial network. After exciting the single target submicron spot, a local response of mitoCa^2+^ rise was activated and generally confined in a subnetwork of mitochondrial tubulars (Fig. 1 B). The surrounding mitochondria and the whole cell remained unperturbed. The photoexcitation efficiency was dependent on both laser power and stimulation duration (Fig. 1 C).

**Figure 1.**
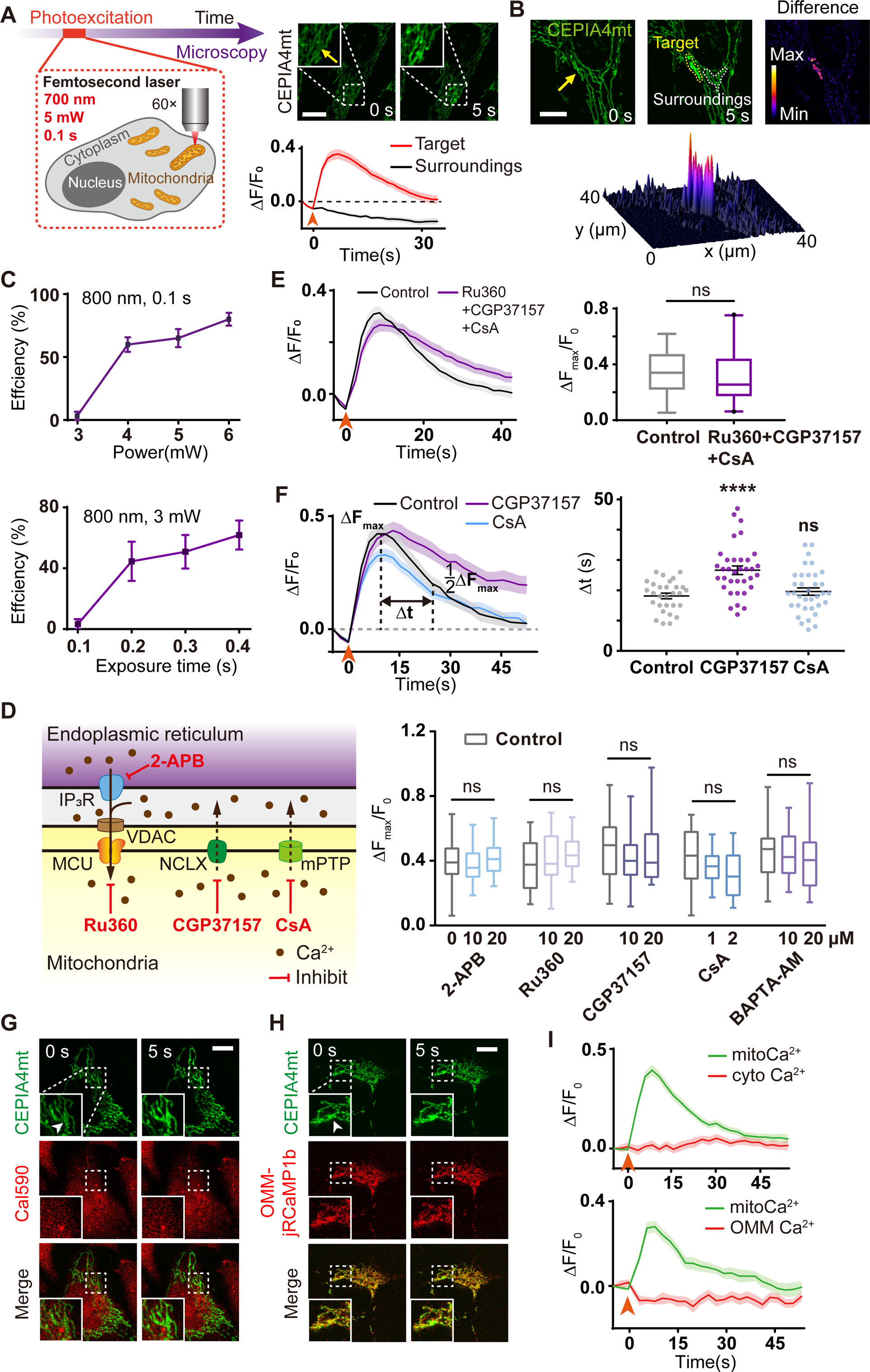
Photoexcitation of an individual mitochondrion triggers local mitoCa^2+^. **(A)** Single-mitochondrial tubular photoexcitation by a tightly-focused femtosecond laser which enables simultaneous and continuous microscopy. Right panel: a photoexcited local mitoCa^2+^ rise indicated by CEPIA4mt (n = 20 cells). Yellow arrow: photoexcitation region. Arrow heads: photoexcitation events. Bar: 10 μm. **(B)** Local mitoCa^2+^ rise restricted in a subnetwork of mitochondria without perturbation to the surroundings. Bar: 10 μm. **(C)** Photoexcitation efficiency is dependent on laser power and exposure time (n = 6 independent groups with 10-20 cells per group) at 800 nm. **(D)** Quantified photoexcited mitoCa^2+^ amplitudes when the mitochondrial ion channels (the channels and their blockers are presented in the left panel) are blocked with 2-APB (n = 20 at 10 μM, n = 20 at 20 μM), Ru360 (n = 34 at 10 μM, n = 16 at 20 μM), CGP37157 (n = 35 at 10 μM, n = 22 at 20 μM) and CsA (n = 34 at 1 μM, n = 16 at 2 μM), respectively. BAPTA-AM (n = 35 at 10 μM, n = 20 at 20 μM) is used to chelate all free cellular Ca^2+^. The control groups are indicated by the 0-concentration. **(E)** Local mitoCa^2+^ transients by photoexcitation simultaneously in the presence with 10 μM Ru360, 10 μM CGP37157 and 1 μM CsA (n = 41 cells in 3 independent experiments). Right panel: quantified mitoCa^2+^ amplitudes. **(F)** MitoCa^2+^ kinetics after photoexcitation in the presence of 20 μM CGP37157 (n = 18 cells in 3 independent experiments) and 2 μM CsA (n = 16 cells in 3 independent experiments). Right panel: quantified decay time △t (the width at half maximum of photoexcited mitoCa^2+^) of mitoCa^2+^ from the left (n = 35 cells in 3 independent experiments). **(G and H)** Simultaneous indication and microscopy of mitoCa^2+^ (indicated by CEPIA4mt) and **(G)** cytosolic Ca^2+^ (indicated by Cal590) or **(H)** Ca^2+^ of outer membrane of mitochondria (indicated by OMM-jRCaMP1b) during photoexcitation. Arrow heads: photoexcitation region. Bar: 10 μm. **(I)** MitoCa^2+^, cytosolic Ca^2+^ (n = 25 cells) and OMM Ca^2+^ (n = 13 cells) kinetics after photoexcitation. \**P* < 0.05, \*\**P* < 0.01, \*\*\**P* < 0.001, \*\*\*\**P* < 0.0001, by two-tailed unpaired t test.

The photoexcited mitoCa^2+^ transients were not involved in any extra-mitochondrial Ca^2+^. To prove this, we tested whether the cytosolic Ca^2+^ influx contributed to the photoexcited mitoCa^2+^ by blocking the Ca^2+^ channels in mitochondria. At first, the endoplasmic reticulum (ER)-mitochondria Ca^2+^ transport was shut down by blocking the Ca^2+^ exit of ER, IP_3_R, and the Ca^2+^ entrance to mitochondria, MCU, separately (Fig. 1 D). The photoexcited mitoCa^2+^ did not show any change. Then two Ca^2+^ exits of mitochondria, NCLX and mPTP were blocked, which did not influence the photoexcited mitoCa^2+^ either (Fig. 1 D). At last, we chelated all free cellular Ca^2+^ but the laser could still trigger consistent mitoCa^2+^ transients. To further confirm this result, mitochondrial Ca^2+^ channels including MCU, NCLX, and mPTP were blocked simultaneously. The local mitoCa^2+^ transients by photoexcitation still stayed the same (Fig. 1 E). Those results suggest the photoexcited mitoCa^2+^ transients were not related to Ca^2+^ transportations from cytosol or ER.

We found the NCLX partially contributed to the discharge of mitoCa^2+^ transients. If NCLX was blocked, the decay speed of mitoCa^2+^ was significantly slower than it of control (Fig. 1 F). The mPTP did not contribute to the mitoCa^2+^ exportation (Fig. 1 F). A slight Ca^2+^ rise in cytosol in the laser spot after photoexcitation could be found (Fig. 1, G and I). The Ca^2+^ level in outer membrane of mitochondria (OMM) did not increase (Fig. 1, H and I). Therefore, mitoCa^2+^ transients could exclusively be formed by local mitochondrial Ca^2+^ stores. This local photoexcitation thus enables us to study the mitoCa^2+^ under a physiological condition that the cellular Ca^2+^ homeostasis maintained during the manipulation of target mitoCa^2+^.

We then clarified how the laser excited mitoCa^2+^ transients. It could be found the photoexcitation efficiency was dependent on wavelength (Fig. 2 A). The spectrum was quite similar to the two-photon absorption spectrum of flavin molecules (Huang et al., 2002). The autofluorescence (490-520 nm) of oxidized flavoproteins (protein-bounded FMN or FAD) consistently colocalized with mitochondria (Fig. 2 B). In this regard, we suspected the photoexcited mitoCa^2+^ transients were mediated by two-photon excitation of the mitochondrial flavoproteins, including complex I (with FMN subunit) and complex II (with FAD subunit) in ETC.

**Figure 2.**
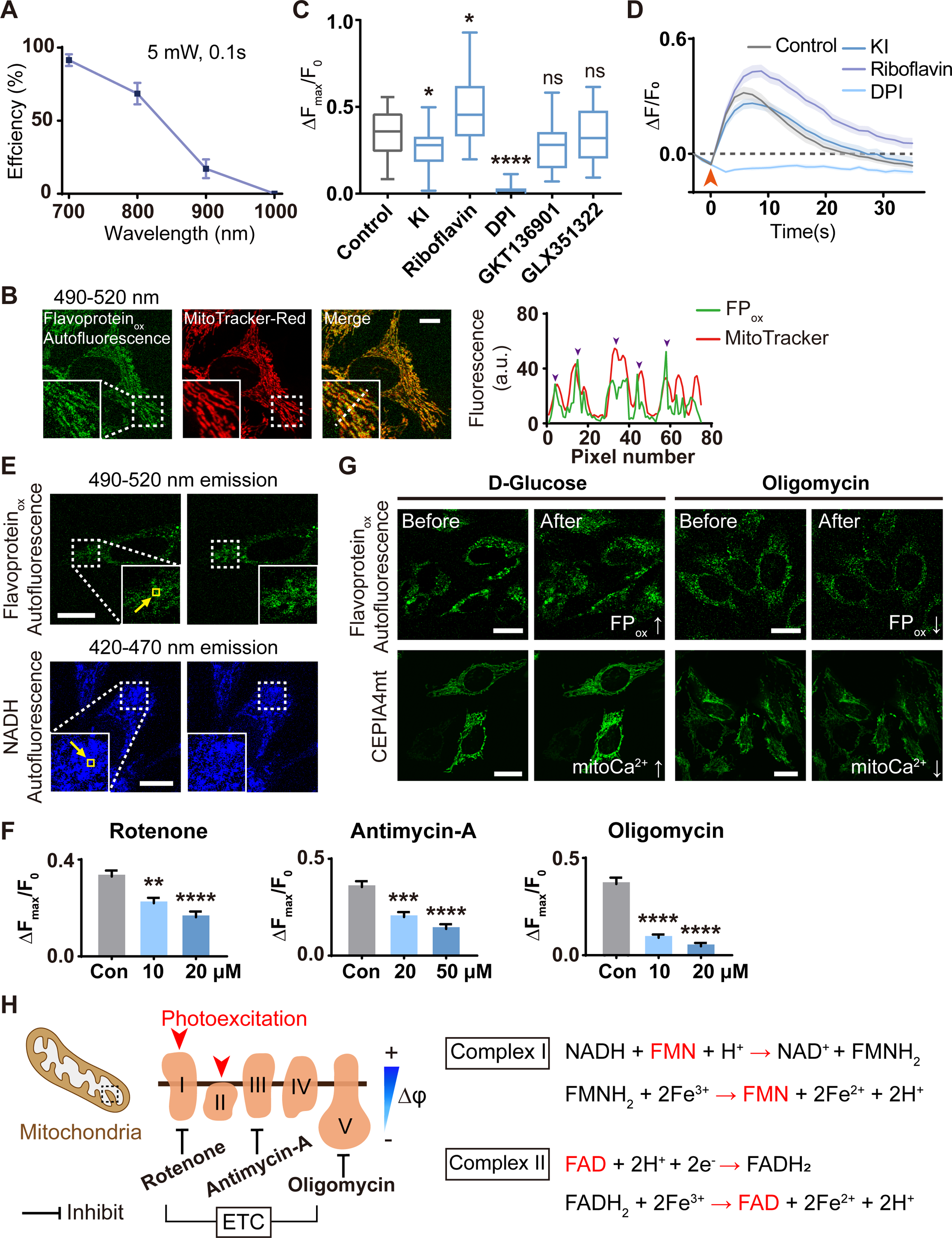
Photochemical excitation to complex I and II accelerates ETC to trigger mitoCa^2+^ transients. **(A)** Spectrum of photoexcitation efficiency (at 5 mW for 0.1 s). n = 7 independent groups of cells with 10-20 cells per group at each point. **(B)** Autofluorescence of oxidized flavoproteins (FP_ox_, excited by 473 nm laser and detected at 490-520 nm) colocalized with MitoTracker Red. Bar: 10 μm. **(C)** Quantified photoexcited mitoCa^2+^ amplitudes in the presence with KI (50 mM, n = 20 cells), riboflavin (10 μM, n = 26 cells), DPI (10 μM, n = 26 cells), GKT136901 (10μM, n = 20 cells) and GLX351322 (10μM, n = 20 cells), respectively. **(D)** MitoCa^2+^ kinetics after photoexcitation in the presence of KI, riboflavin and DPI. Arrow heads: photoexcitation events. **(E)** Autofluorescence of FP_ox_ (n = 10 cells) and NADH (excited by 720 nm femtosecond laser and detected at 420-470 nm, n = 11 cells) before and after photoexcitation respectively. Yellow arrows: photoexcitation region. Bar: 10 μm. **(F)** Photoexcited amplitudes of mitoCa^2+^ in the presence with Rotenone (n = 43 cells at 10 μM, n = 36 cells at 20 μM), Antimycin-A (n = 38 cells at 20 μM, n = 20 cells at 50 μM) and DPI (n = 40 cells at 10 μM, n = 32 cells at 20 μM), respectively. **(G)** Autofluorescence of FP_ox_ and mitoCa^2+^ before and after D-Glucose or oligomycin treatment, respectively. Bar: 10 μm. **(H)** Acceleration of electron transport at complex I and II by photoexcitation to flavoproteins there. Red words (FMN and FAD): the photoexcited local flavoproteins. \**P* < 0.05, \*\**P* < 0.01, \*\*\**P* < 0.001, \*\*\*\**P* < 0.0001, by two-tailed unpaired t test.

To check this hypothesis, we photoexcited mitochondria while inhibiting/quenching photoexcited free flavin, protein-bound flavin (generally complex I and II), NADPH oxidase (NOX) 1/4 and NOX 4, separately (Fig. 2 C). Only the inhibition of flavoproteins suppressed the photoexcited mitoCa^2+^ significantly, indicating the photoexcitation target was complex I and II to produce mitoCa^2+^ transients (Fig. 2, C and D). Free flavin molecules and NOX contributed little in this process. The photoexcited mitoCa^2+^ amplitude increased significantly if upregulating cellular flavin concentration by adding riboflavin (RF) to the cell medium for one-day culture, consistent with this result (Fig. 2, C and D).

After photoexcitation, the autofluorescence of oxidized flavoproteins increased while the NADH fluorescence decayed accordingly (Fig. 2 E), implying the acceleration of NADH dehydrogenation and the oxidization of FMNH_2_ to transfer electrons to ubiquinone (coenzyme Q). To examine if the photoexcitation of mitoCa^2+^ relied on the entire ETC, we inhibited complex I, III, and ATP synthase (complex V) separately and found the photoexcited mitoCa^2+^ was significantly suppressed in each case (Fig. 2 F). Consistently, if the cells were treated with D-Glucose, both the autofluorescence of flavoproteins and mitoCa^2+^ increased significantly (Fig. 2 G). If the ATP synthesis was inhibited, the autofluorescence of flavoproteins and mitoCa^2+^ both decreased significantly (Fig. 2 G). Those results suggest the mitoCa^2+^ transients were generated by the acceleration of electron transports at Complex I and II by photoexcitation to flavoproteins there (or directly by glucose) (Fig. 2 H).

### 2. ATP synthesis releases Ca^2+^ to mitochondrial matrix

We then specified how mitoCa^2+^ was finally released in this photoexcitation process. At first, the photoexcitation induced a transient significant Mg^2+^ decay in the targeted mitochondrial region, suggesting the simultaneous enhanced ATP synthesis (Fig. 3 A). We then constructed ATP5B siRNA to knock down ATP5B (the β subunit of ATP synthase), verified by immunoblotting (Fig. 3 B), to suppress ATP synthesis (von Ballmoos et al., 2009). The knockdown (KD) of ATP5B was further verified by using immunofluorescence microscopy of it to confirm the lower level and localization of ATP5B in mitochondria (Fig. 3 C). In those ATP5B-KD cells, the photoexcited mitoCa^2+^ transients were significantly weaker than them in control (Fig. 3 D). Their decay was also faster (Fig. 3 E). Those results imply mitoCa^2+^ was released during ATP synthesis.

**Figure 3.**
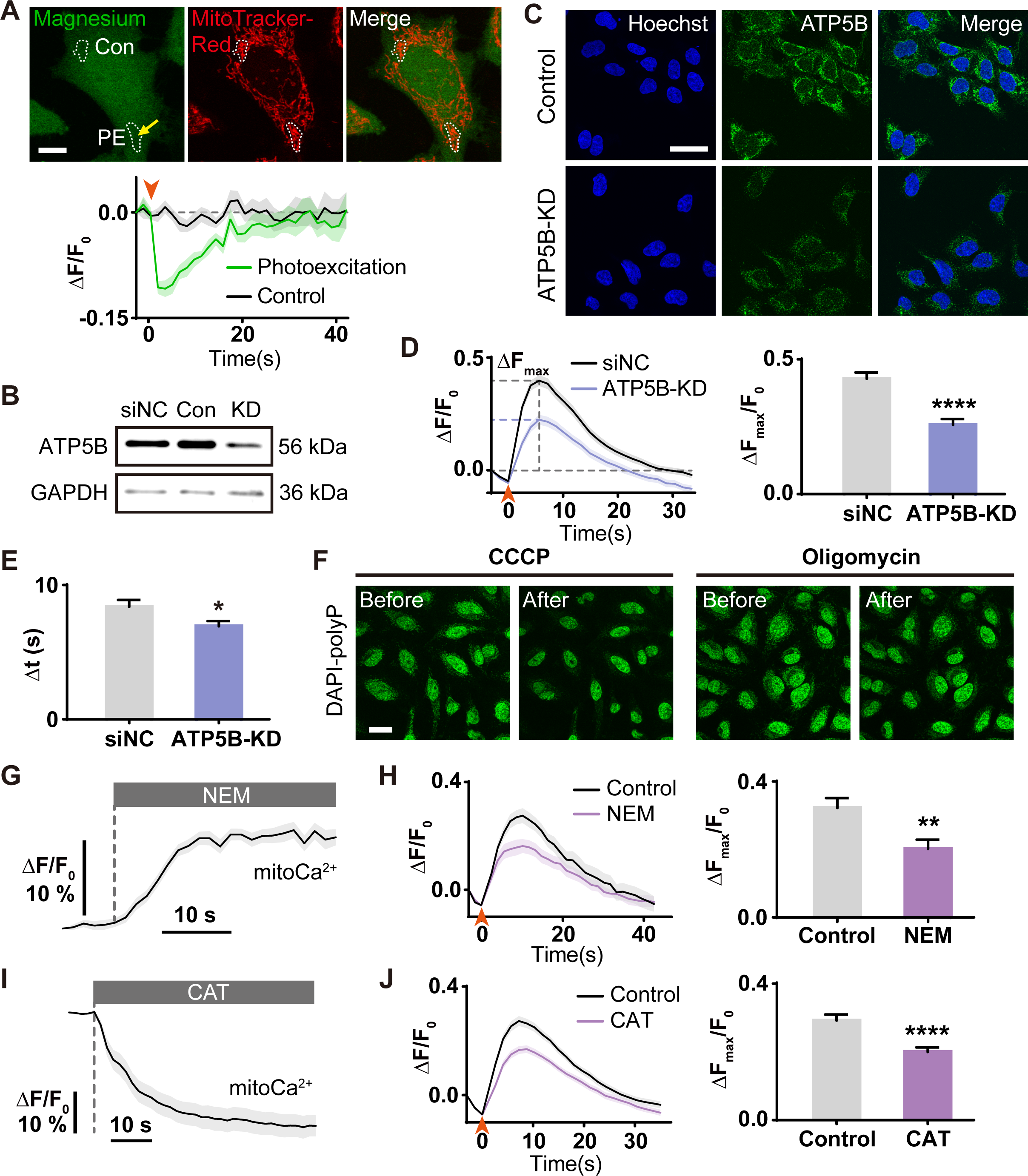
ATP synthesis releases mitoCa^2+^. **(A)** Cells stained with Magnesium Green and MitoTracker Red. Lower panel: a temporary decrease of Mg^2+^ in the photoexcited mitochondria (n = 8 cells, 3 trials). Con: control. PE: photoexcitation. Yellow arrow: photoexcitation region. Arrow heads: photoexcitation events. Bar: 10 μm. **(B)** Western blot of ATP5B in negative control (siNC, transfected with an invalid siRNA), control and ATP5B-KD HEK293 cells. **(C)** Immunofluorescence microscopy of ATP5B in control and ATP5B-KD HEK293 cells. Bar: 50 μm. **(D)** Photoexcited mitoCa^2+^ transients of ATP5B-KD cells (n = 45 cells in 5 independent experiments). Right panel: quantified mitoCa^2+^ amplitudes from those transients. **(E)** Decay time △t of mitoCa^2+^ in ATP5B-KD cells (n = 35 cells in 5 independent experiments). **(F)** Poly-phosphate (polyP) ions in mitochondria indicated by DAPI before and after CCCP or oligomycin treatment. Bar: 30 μm. **(G)** Acute mitoCa^2+^ response to NEM treatment (n = 26 cells in 3 independent experiments). **(H)** Photoexcited mitoCa^2+^ kinetics of cells treated with 10 μM NEM for 30min (n = 23 cells in 3 independent experiments). Right panel: quantified mitoCa^2+^ amplitudes from the left. **(I)** MitoCa^2+^ response to CAT treatment (n = 20 cells in 3 independent experiments). **(J)** Photoexcited mitoCa^2+^ kinetics in cells treated with 100 μM CAT for 1 min (n = 31 cells in 3 independent experiments). \**P* < 0.05, \*\**P* < 0.01, \*\*\**P* < 0.001, \*\*\*\**P* < 0.0001, by two-tailed unpaired t test.

To examine this hypothesis, we manipulated the substrates of ATP synthesis to study the photoexcited mitoCa^2+^ transients. At first, mitochondrial poly-phosphate (polyP) ions were indicated by DAPI by forming DAPI-polyP complexes (Aschar-Sobbi et al., 2008), which decreased after CCCP and oligomycin treatment (Fig. 3 F). If the transport of Pi into mitochondria was blocked, immediately mitoCa^2+^ increased (Fig. 3 G). Under this condition, the photoexcited mitoCa^2+^ was suppressed (Fig. 3 H). Those results suggest that Ca^2+^ buffered in mitochondria partially exists in the form of Ca^2+^-Pi complexes, consistent with the previous theory (Chalmers and Nicholls, 2003). If the supply of ADP was blocked, an acute decline of mitoCa^2+^ could be found (Fig. 3 I). After that, the amplitude of mitoCa^2+^ transients trigged by photoexcitation was significantly lower than it of control (Fig. 3 J). Taken together, those results suggest the photoexcited mitoCa^2+^ was released from the Ca^2+^-Pi complexes during ATP synthesis.

We then demonstrated the transient acceleration of ETC by photoexcitation depleted H^+^ for ATP synthesis and subsequently induced depolarization of mitochondrial membrane potential (MMP), which eventually suppressed mitoCa^2+^ in turn. In the experiments, we found the mitochondria exhibited photoexcited mitoCa^2+^ transients were depolarized (Fig. 4 A). Such MMP depolarization was not caused by the photoexcited mitoCa^2+^ rise, verified by introducing global Ca^2+^ rise of the whole cell and remarkable mitoCa^2+^ influx that did not induce immediate MMP depolarization (Fig. 4 B). Meanwhile, the mitochondrial pH (mito-pH) increased in both mitochondrial matrix and inner membrane space (IMS) after photoexcitation, suggesting the depletion of H^+^ (Fig. 4 C). Here the mitochondrial pH increase did not influence the fluorescence of mitoCa^2+^ indicator CEPIA4mt (Fig. 4 D).

**Figure 4.**
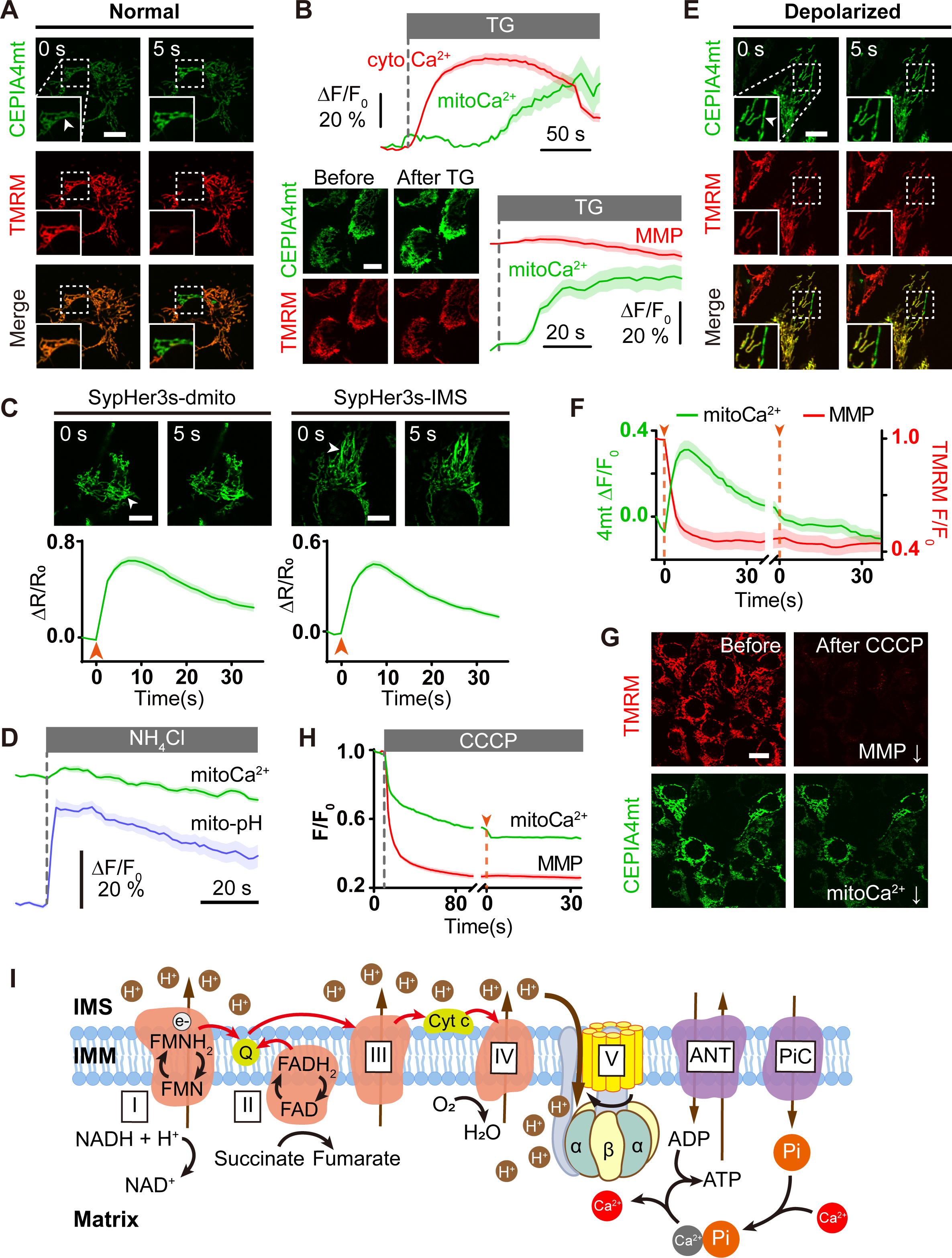
Photoexcited ETC acceleration depletes H^+^ for ATP synthesis and mitoCa^2+^ release. **(A)** Photoexcitation of mitochondria induced local mitoCa^2+^ rise (indicated by CEPIA4mt) and MMP depolarization (indicated by TMRM). Bar: 10 μm. **(B)** Sole mitoCa^2+^ rise does not lead to MMP depolarization, tested by 10 μM TG treatment (n = 13 cells in 3 independent experiments). Bar: 10 μm. **(C)** PH level in mitochondrial matrix and IMS after photoexcitation were indicated by SypHer3s-dmito and SypHer3s-IMS respectively. (n =31 and 43 cells in 3 independent experiments respectively). Arrow heads: photoexcitation events. Bar: 10 μm. **(D)** Verification of the mitoCa^2+^ indicator, CEPIA4mt, against the mitochondrial pH increase, tested by 5 mM NH_4_Cl treatment to introduce alkalinity (n = 18 cells in 3 independent experiments). **(E)** Photoexcitation can not trigger any mitoCa^2+^ responses in depolarized mitochondria (n = 20 cells). Bar: 10 μm. **(F)** MitoCa^2+^ responses to the second photoexcitation after MMP depolarization by the first one (n = 17 cells in 3 independent experiments). **(G)** MMP depolarization by 10 μM CCCP directly leads to a mitoCa^2+^ decrease (n = 5 independent experiments). Bar: 20 μm. **(H)** MitoCa^2+^ responses to photoexcitation in the presence of CCCP (n = 27 cells in 3 independent experiments). **(I)** Proposed mechanism of mitoCa^2+^ release from Ca^2+^-Pi complexes through ATP synthesis. Red arrows indicate the transfer of electrons.

In reverse, if the mitochondria were depolarized by photoexcitation at first, the ETC could not work, and then the laser could not activate any mitoCa^2+^ (Fig. 4, E and F). To confirm this, we used CCCP to depolarize mitochondria and found simultaneous mitoCa^2+^ decline (Fig. 4 G), consistent with our theory. In this case, if the mitochondria suffered photoexcitation, no mitoCa^2+^ transients could be triggered (Fig. 4 H). Hence it should be the photoexcited large H^+^ current at ATP synthase that took the responsibility of MMP depolarization. According to this photochemical excitation and the mitoCa^2+^ transients, we established such a model that the mitoCa^2+^ transients require MMP for maintaining ETC, which are released from the Ca^2+^-Pi complexes at complex V through ATP synthesis driven by the excess H^+^ current accumulated by ETC (Fig. 4 I).

### 3. Low mitoCa^2+^ leads low proliferation and low death

We finally demonstrated the knockdown (KD) of ATP synthase subunit that directly influenced the mitoCa^2+^ release led low life activities, by constructing the ATP5B-KD cells. ATP5B-KD cells presented significantly lower proliferation rate than the control (Fig. 5, A-C). However, those ATP5B-KD cells also exhibited death resistance under high oxidative stress, probably due to the lower mitoCa^2+^ level that prevented cells from initiation of apoptosis pathways (Fig. 5 D). A more direct presentation of the ATP synthase influence on mitoCa^2+^ was the recruitment of Parkin induced by CCCP treatment. Cohen’s d values of Parkin immunofluorescence intensity for control (1.54) and ATP5B-KD cells (1.18) suggest mitophagy was more difficult to initiate in ATP5B-KD cells (Fig. 5 E). We finally compared the autophagy level of ATP5B-KD cells with control that were treated by CCCP and found the LC3 clusters in ATP5B-KD cells were significantly fewer than them in control (Fig. 5 F). Therefore, those ATP5B-KD cells showed resistance to autophagy (mitophagy) and death, but also with low proliferation ability.

**Figure 5.**
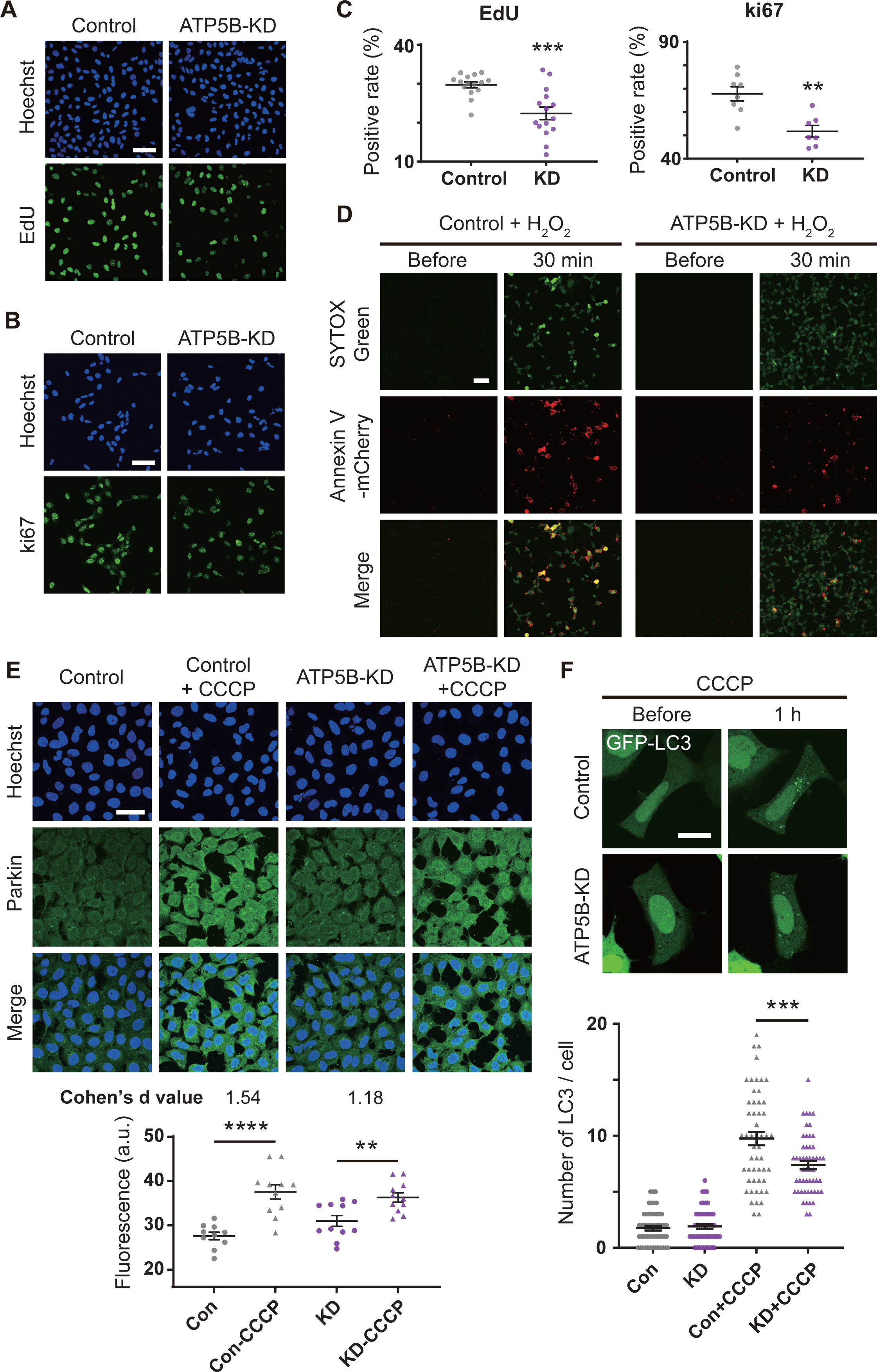
Phenotypes related to mitoCa^2+^ level of ATP5B-KD cells. **(A)** EdU test of ATP5B-KD cells. Bar: 80 μm. **(B)** Immunofluorescence images of ki67 in ATP5B-KD cells. Bar: 80 μm. **(C)** Quantified ratio of EdU- and ki67-positive ATP5B-KD cells (n = 7 fields and 15 fields in 3 independent experiments, respectively). **(D)** SYTOX Green and Annexin V-mCherry test of ATP5B-KD cells before and after 5 mM H_2_O_2_ treatment for 30 min. Bar: 80 μm. **(E)** Immunofluorescence imaging of Parkin in ATP5B-KD cells before and after 10 μM CCCP treatment for 1 h (replicated three times). Lower panel: quantified fluorescence intensity of Parkin (n = 9 fields) and the calculated Cohen’s d values accordingly. Bar: 50 μm. **(F)** The statistics of LC3 clusters in control and ATP5B-KD cells before and after 1-h CCCP (10 μM) treatment, respectively. Lower panel: number of LC3 clusters per cell (n = 52 cells). Bar: 20 μm. \**P* < 0.05, \*\**P* < 0.01, \*\*\**P* < 0.001, \*\*\*\**P* < 0.0001 by two-tailed unpaired t test.

## Discussion

Mitochondria have long been recognized as Ca^2+^ buffer to coordinate intracellular Ca^2+^ homeostasis. Therefore, extra-mitochondrial Ca^2+^, particularly the cytosolic Ca^2+^, is believed as the main source to dominate mitoCa^2+^ and further mitochondrial signaling pathways, functions, and metabolism through influx into mitochondria, instead of local mitoCa^2+^ (Boyman et al., 2021; Loncke et al., 2021). The view that mitoCa^2+^ regulates respiration largely relies on the change of metabolism substrates. A series of previous works suggest the Ca^2+^ influx into mitochondria promotes respiration, probably by Ca^2+^-sensitive dehydrogenases or even interaction with complex V (Denton, 2009; Glancy and Balaban, 2012; Wescott et al., 2019). The evidence is generally the curve of consume of ATP and Ca^2+^ input to mitochondria (Wescott et al., 2019). However, the causality between mitoCa^2+^ and respiration is still elusive, which is quite challenging to clarify since those two factors can be the cause and consequence of each other. In cells, if cytosolic Ca^2+^ enters mitochondria that elevates mitoCa^2+^, the cellular Ca^2+^ has already been abnormal. In this case, the cellular Ca^2+^ pumps, like PMCA and SERCA, start to pump cytosolic Ca^2+^ into ER or out of the cell, which all burn off ATP and mitochondria need to accelerate respiration to supply such energy (Bruce, 2018; Periasamy et al., 2017). During the respiration, mitochondria release more Ca^2+^ by ATP synthesis to form a loop. This caused a very high correlation between Ca^2+^ and respiration. Therefore, the detection of respiration substrates could hardly specify the mitoCa^2+^ source and its relationship with respiration. According to our data, high respiration generates more mitoCa^2+^, rather than the reverse.

In this study, we report a phenomenon of photoexcited mitoCa^2+^ transients in which the two-photon excitation to complex I and II accelerates ETC to release mitoCa^2+^. The specificity is guaranteed by the precision of the laser focus onto mitochondrial tubulars and the specific two-photon absorption of local flavoproteins. A blue laser (460 ∼ 490 nm) was not used here to directly excite flavoproteins because the unspecific absorption by other molecules to the blue laser with high-energy photons can hardly be avoided. The direct photodamage and oxidative stress by the blue lasers also introduce more perturbations to mitochondria and cells.

The point that free Ca^2+^ in mitochondrial matrix forms complexes with Pi spontaneously has been reported previously as a potential form of Ca^2+^ existence in mitochondria (Chalmers and Nicholls, 2003; Duvvuri and Lood, 2021; Solesio et al., 2016; Wei et al., 2015). When ADP and Pi are consumed to produce ATP, Ca^2+^ is then released from the Ca^2+^-Pi complexes. The free mitoCa^2+^ can re-combine with Pi to decrease the free mitoCa^2+^ level. This machinery maintains a physiological level of mitoCa^2+^ in a simple manner. Such local mitoCa^2+^ release does not induce the opening of mPTP while is slowly exported through NCLX. Although it is difficult to directly observe the Ca^2+^-Pi complexes, the data of mitoCa^2+^, ATP synthase, Pi (and polyP), and ADP are all consistent with this theory. Our results thus present a deep insight to the mitoCa^2+^ release and regulation machinery.

## Author contributions

H.H. and Y.Z. conceived the study and supervised the project. B.L. performed the experiments and prepared the figures. H.H. drafted the manuscript. All authors analyzed the data, discussed the results, and revised the manuscript.

## Acknowledgement

We thank Bei Ding for the constructive discussions. This work was supported by the National Natural Science Foundation of China (NSFC 61975118, 62205197, 92054105, and 62022056), and Shanghai Jiao Tong University (YG2022QN063).

## Data and Materials Availability

The data that support the findings of this study are available from the corresponding author upon reasonable request.

## Methods

### 1. Cells and materials

Hela and HEK293 cells were cultured in Dulbecco’s Modified Eagle’s Medium (DMEM, Gibco) supplemented with 10% fetal bovine serum (FBS, Gibco) and 1% penicillin/streptomycin (Gibco). All cells were incubated in 90-mm dishes (Corning) and 35-mm glass bottom dishes (Biosharp) at 37 ℃ in humidified 5% CO_2_ atmosphere.

For the inhibitor experiments, potassium iodide (KI) was purchased from Sigma-Aldrich. Diphenyleneiodonium chloride (DPI), GKT136901, GLX351322, rotenone, oligomycin, Ru360, 2-Aminoethyl diphenylborinate (2-APB), cyclosporin A (CsA), CGP37157, N-Ethylmaleimide (NEM), and carboxyatractyloside (CAT) were purchased from MedChemExpress (MCE). Ru360 was purchased from Santa Cruz. BAPTA-AM was purchased from ThermoFisher. Antimycin-A was purchased from MKBio. KI, DPI, GKT136901 and GLX351322 were used to inhibit photoexcited flavin, protein-bound flavin, NADPH oxidase (NOX) 1/4 and NOX 4, respectively. Rotenone, antimycin-A and oligomycin were used to inhibit complex I, complex III and ATP synthase (complex V), respectively. 2-APB, Ru360, CGP37157 and CsA were used to block Ca^2+^ channels IP_3_R, MCU, NCLX and mPTP, respectively. NEM and CAT were used to inhibit mitochondrial phosphate carrier and ADP/ATP carrier. To clear the intracellular Ca^2+^, cells were incubated in Ca^2+^ free buffer with 10μM BAPTA-AM at 37 ℃ for 30min. The Ca^2+^ free buffer contained 140 mM NaCl, 5 mM KCl, 1 mM MgCl_2_ (all from Sinopharm Chemical Reagent), 10 mM D-glucose (BBI Life Sciences), 10 mM HEPES (Gibco), and 10 μM EGTA (Sigma), whose pH was titrated to 7.4 with NaOH. Cells were incubated in the medium with corresponding inhibitors for 30-60 min before conducting photoexcitation. Other chemicals included riboflavin (Sigma), NH_4_Cl (Macklin) and CCCP (Sigma). Cells were incubated in the medium added with 10μM riboflavin (RF) for 24 h before photoexcitation. Cells were treated with 5 mM NH_4_Cl to introduce alkalinity. 10 μM CCCP was used to treat cells for 1 h to induce autophagy (mitophagy).

Fluorescent dyes including MitoTracker Red (100 nM, ThermoFisher), TMRM (100 nM, ThermoFisher), Magnesium green^TM^ AM (4 μM, ThermoFisher) and Cal590 (5 μM, AAT Bioquest) were used for the indication of mitochondria, MMP, cellular Mg^2+^ and cytosolic Ca^2+^, with which cells were incubated for 15 min, 15 min, 30 min and 1 h, respectively. Then the medium was replaced with fresh DMEM before fluorescence imaging. For mitochondrial polyP imaging, cells were incubated with DAPI (5 μg/ml, Sigma) for 14 h.

### 2. Plasmids construction and transfection

CEPIA4mt (Plasmid #58220), SypHer3s-dmito (Plasmid #108119), SypHer3s-IMS (Plasmid #108120), OMM-jRCaMP1b (Plasmid #127871) and GFP-LC3 (Plasmid #22405) were purchased from Addgene, transfected to the cells for the indication of mitoCa^2+^, mitopH, pH of IMS, Ca^2+^ of outer membrane of mitochondria and LC3, respectively. Cells were passaged in 35-mm glass bottom dishes for 24h incubation then transfected with 1ug plasmid per dish for 4-5 h using jetPRIME^®^ transfection reagent (Polyplus). Before conducting fluorescence microscopy, the transfected cells were incubated for 20 h with DMEM medium replaced.

### 3. Knockdown of ATP5B and Western blot

ATP5B (NCBI Gene ID 506) was knocked down by siRNA. The sense sequence was 5’-GUAUGGAUGAACUUUCUGA(dT)(dT)-3’. HEK293 cells were transfected with 50 nM siRNA for 20 h using jetPRIME ^®^ transfection reagent.

The expression of ATP5B in knockdown cells was verified by Western blotting (WB). Cells were dissected in lysis buffer (RIPA: protease inhibitor = 100: 1) on the ice for 15 min. Then the mixture was centrifuged to extract proteins for subsequent western blotting. Proteins were electrophoretic separated on 4-20% SDS-PAGE precast gels (Beyotime) and transferred to a Hybond P polyvinylidene difluoride membrane (Millipore). The membrane was blocked in 5% skim milk for 2 h at room temperature then incubated with primary antibodies (Rabbit monoclonal to ATPB at 1/50000 dilution, Abcam) for 16 h at 4 ℃, followed by incubation with secondary antibodies (1/10000 dilution, Abcam) for 2 h at room temperature. Proteins were visualized using BeyoECL Star kit (Byotime). For detection of internal reference GAPDH, the membrane was soaked with western blot fast stripping buffer (EpiZyme) to remove the primary antibodies of anti-ATPB, then re-blocked and conducted with subsequent WB operations.

### 4. Microscopy

All fluorescence imaging was performed using confocal microscope (FV1200, Olympus). ×20 (N.A. = 0.7) objective was used to image the cells stained with SYTOX Green and Annexin V-mCherry. ×30 (oil immersion, N.A. = 1.05) objective was used for the imaging in Edu test and ki67 immunofluorescence microscopy. Other imaging was performed with ×60 (water immersion, N.A. = 1.2) objective for finer microscopy. CEPIA4mt and magnesium green were excited by 473 nm laser and detected by photomultiplier tube (PMT) with a 490-520 nm filter before it. OMM-jRCaMP1b, MitoTracker-Red, TMRM and Cal590 were excited by 543 nm laser and detected at 560-620 nm. Images were captured at a speed of 2 μs per pixel. Confocal images of flavoprotein autofluorescence excited by 473 nm laser and detected at 490-520 nm were captured with 1024 × 1024 pixels per frame at 2 μs per pixel. NADH autofluorescence was two-photon excited by 720 nm femtosecond laser and detected at 420-470 nm. Ten frames were superposed to acquire an average image of flavoproteins or NADH autofluorescence with high signal-to-noise ratio. DAPI-polyP complex was excited by 405 nm laser and detected at 490-520 nm.

### 5. Immunofluorescent microscopy

The confocal microscope was used for immunofluorescent microscopy. Cells were fixed with 5% paraformaldehyde and permeabilized with 0.3% Triton X-100, then blocked in 5% skim milk for 1 h at room temperature. The cells were incubated with primary antibodies (Rabbit monoclonal to ATPB at 1/100 dilution, Abcam) for 14 h at 4 ℃, followed by incubation with secondary antibodies (Alexa Fluor 488 at 1/1000 dilution, Abcam) for 1 h at room temperature. Ki67 rabbit antibody (1/100 dilution, Abcam) and Parkin rabbit antibody (1/50 dilution, ThermoFisher) were used according to the protocols as well. Cell nuclei were stained using Hoechst 33342 (Byotime) excited by 405 nm laser.

### 6. Photoexcitation to individual mitochondria

The photoexcitation method was established based on a confocal microscope coupled with a femtosecond laser (tunable from 680-1080 nm, 140 ± 20 fs, 80 MHz, Chameleon Ulttra II, Coherent). The femtosecond laser was controlled by a mechanical shutter, collimated to the galvo-mirrors in the microscope, and finally tightly focused by a 60 × (water immersion, N.A. = 1.2) objective. In this way, the femtosecond-laser focus could be located in any predefined target by controlling the galvo-mirrors. In experiments, cells were passaged in 35-mm glass bottom dishes (Biosharp) before photoexcitation. The photoexcitation was localized to individual mitochondria by selecting the target as the field of interest and defining the photoexcitation as a single frame, which could be inserted at any predefined time slot in a continuous time-lapse confocal microscopy sequence. When the microscopy task started, the shutter was open synchronized with the photoexcitation frame to enable the femtosecond laser focus onto the target mitochondrial tubular for a predefined duration (the frame time). After that, the shutter closed and the time-lapse microscopy continuously provided the fluorescent images of the cells.

### 7. Data processing and statistics

The data processing was completed using GraphPad Prism 7. The error bars displayed in graphs represented mean ± SEM. Significance test was analyzed by two-tailed unpaired t-test. Box plots were plotted with 2.5-97.5 percentile. ImageJ was used for fluorescence intensity statistics. Fluorescence from the nucleus was excluded in Parkin immunofluorescence intensity statistics. Cohen’s d was calculated by (μ_1_-μ_2_)/ σ, where (μ_1_-μ_2_) and σ are the mean difference of Parkin immunofluorescence intensity before and after CCCP treatment and standard deviation, respectively.

